# Monocyte-derived macrophage recruitment mediated by TRPV1 is required for eardrum wound healing

**DOI:** 10.1101/2025.02.03.635565

**Authors:** Yunpei Zhang, Pingting Wang, Lingling Neng, Kushal Sharma, Allan Kachelmeier, Xiaorui Shi

## Abstract

The tympanic membrane (TM), or eardrum, is a thin, sensitive tissue critical for hearing by vibrating and transmitting sound waves to the inner ear. TM perforation and development of otitis media and conductive hearing loss are commonly seen in the clinic. In this study, we demonstrate the role of TRPV1 signaling mediated macrophage recruitment and angiogenesis in TM repair. By creating a wounded TM mouse model with a perforation in the anteroinferior region of the pars tensa — a region in humans often damaged in traumatic injury, we observed a massive accumulation of macrophages in the vicinity of the acutely wounded TM. Using 5-Ethynyl-2’-deoxyuridine pause labeling and a chimeric bone marrow transplant model, we found that most of the recruited macrophages did not originate from local tissue-resident macrophages but rather from blood-circulating monocytes. Parallel to macrophage recruitment, angiogenesis was observed near the wound on day 3 after perforation and further progressed by day 7. The angiogenic process was strongly associated with the recruited macrophages, as macrophage depletion resulted in a notable reduction in angiogenesis. At the transcriptional level, we found that macrophages facilitate angiogenesis through several signaling pathways. Additionally, we identified direct intercellular communication between macrophages and endothelial cells mediated by phosphoprotein 1 signaling. Furthermore, Gene Ontology analysis of bulk RNA sequencing data from TMs revealed that the macrophage recruitment is associated with neuroinflammatory responses. Using a fluorescence reporter mouse driven by TRPV1, we discovered that the TM contains rich sensory nerve fibers expressing TRPV1. A genetic mutation in the *Trpv1* gene resulted in a marked decrease in the expression of neuroinflammatory genes, such as *Tac1*. This decrease subsequently resulted in reduced macrophage recruitment, impaired angiogenesis, and delayed wound healing. Together, these findings highlight the crucial role of TRPV1 signaling in monocyte migration and macrophage-related angiogenesis, both of which are crucial for facilitating healing of the TM. These results also open new opportunities for clinical interventions. Targeting TRPV1 signaling could enhance TM immunity, improve blood circulation, promote the repair of damaged TM, and ultimately prevent middle ear infections.

## Introduction

The normal structure and function of the tympanic membrane (TM) are essential for sound transduction and hearing. Sound waves are transmitted through the external auditory meatus, causing the TM to vibrate. These vibrations are then transmitted to the auditory ossicles, the small bones in the middle ear, which amplify the sound vibrations before passing them on to the inner ear. Sensory hair cells in the inner ear convert these mechanical signals into electrical signals that the brain interprets as sound. Additionally, the TM serves as a protective barrier, shielding the middle ear from invading pathogens. Anatomically, the TM consists of two regions: the pars tensa and the pars flaccida. The pars tensa is the largest portion of the TM and vibrates in response to sound. The pars flaccida, located in the superior portion of the malleus, helps maintain the proper function of the pars tensa by stabilizing pressure in the middle ear (Stenfors et al. 1979). The pars tensa is a thin and delicate structure that is often vulnerable to injury (Paik et al. 2022; Pusz and Robitschek 2017; Van Hoecke et al. 2016; Littlefield and Brungart 2020). Traumatic perforations, which can occur due to assault or sound blasts, are commonly observed in young adults. Although most TM perforations heal spontaneously, approximately 20% do not, which can lead to complications such as middle ear infections and conductive hearing loss (Santa Maria et al. 2007; Sagiv et al. 2018; Dolhi and Weimer 2020; Lou 2021; Lou et al. 2012).

Unlike typical cutaneous wounds, the TM is unique as a healing tissue because it is suspended in air. During the healing process, reparative cells must bridge a gap without a ready-made surface over which they can migrate (Santa Maria et al. 2010). Previous studies have shown that the squamous epithelium, the outer layer of the TM, is responsible for closing the perforation by forming an epithelial bridge over the lesion (Araujo et al. 2014; Santa Maria et al. 2010; Rosowski et al. 2011; Kitazawa et al. 2015; Bergevin and Olson 2014; Aarnisalo et al. 2010; Kurabi et al. 2017; Henson and Henson 2000; Dong et al. 2019; Frumm et al. 2020). More recent research indicates that the localization of progenitor or stem cells in the epidermal layer plays a significant role in TM repair (Mozaffari et al. 2020; Frumm et al. 2020). A study by Scaria et al. demonstrated that after injury, a multi-lineage blastema-like cellular mass is recruited, which forms a wound epidermis (Scaria et al. 2023). This finding highlights that TM repair is a type of epimorphic regeneration. However, a comprehensive understanding of the complex network of cellular processes and molecular interactions involved in TM wound healing, as well as the mechanisms leading to failed healing, remains limited. In this study, we report that monocyte recruitment mediated by TRPV1 (transient receptor potential vanilloid 1) signaling expressed in peripheral nerve fibers of TM, along with macrophage-associated angiogenesis, are crucial during the acute phase of TM wound healing.

Wound healing is a highly orchestrated and complex regulatory process that involves inflammation, proliferation, and remodeling (Wilkinson and Hardman 2020). Recent studies have highlighted the vital role of macrophages in wound healing, including their functions in dampening inflammation, clearing cellular debris, and coordinating tissue repair (Krzyszczyk et al. 2018; Kim and Nair 2019). Dysfunctional macrophage responses can lead to delayed wound healing and tissue fibrosis (Lech and Anders 2013). In our study, we found that the macrophages recruited to the injured area following TM injury are derived from monocytes, rather than being locally proliferated or migrated from resident tissue macrophages. Alongside the recruitment of monocytes, we also observed the growth of new blood vessels, which is a crucial element in TM repair. The presence of macrophages in the wound area is strongly linked to angiogenic activity, as depleting macrophages significantly impairs angiogenesis. Through the analysis of recent single-cell RNA sequencing datasets (GEO; accession number GSE196692) obtained from mouse TMs at various time points following perforation (Scaria et al. 2023), we found that macrophages promote angiogenesis during the early stages of wound healing via different signaling pathways. Notably, we discovered that communication between macrophages and endothelial cells is mediated by secreted phosphoprotein 1 (*Spp1*).

The TM is densely innervated with sensory nerve fibers (Uddman et al. 1988). Clinically, perforation of the TM can cause a sudden onset of pain in patients due to the injury and stretching of nerves. Using a transgenic TRPV1 fluorescence reporter mouse line, we unexpectedly found that the TM contains a high density of TRPV1-expressing sensory nerve fibers. TRPV1 is a non-selective cation channel and polymodal receptor activated by agents such as capsaicin, endogenous lipids, heat, and mildly acidic pH (Ho et al. 2012; Marrone et al. 2017). It is known for its rapid response to noxious stimuli (Ho et al. 2012; Silverman et al. 2020). Studies have illustrated that the activation of TRPV1 can quickly initiate a cascade of events, including vascular inflammation, immune responses, and proliferation of precursor cells. Research in other organ systems has specifically shown that damage-induced TRPV1 activation triggers a rapid release of neuropeptides, facilitating macrophage recruitment, vascular cell proliferation, and wound healing (Feng et al. 2017; Foster et al. 2017; Wang et al. 2020; Bujak et al. 2019; Schäffer et al. 1998; Ziche et al. 1990; Yano et al. 1989; Nguyen et al. 1995). In this study, we demonstrate for the first time the role of TRPV1 signaling in macrophage mobilization and macrophage-related angiogenesis, which contributes to the healing of TMs, as *Trpv1^-/-^* mice exhibited delayed and impaired wound healing with reduced monocyte recruitment. Our findings suggest that targeting TRPV1 signaling and the associated recruitment of monocytes may be an effective strategy for restoring damaged TMs by enhancing blood circulation — an essential factor often lacking in failed TM restoration efforts.

## Materials and methods

### Animals

The mouse strains used in this study include C57BL/6J (stock# 000664), CSF1R*^EGFP^* (B6.Cg-Tg(Csf1r-EGFP)1Hume/J, stock# 018549), CX3CR1*^CreER^* (B6.129P2(Cg)-*Cx3cr1^tm2.1(cre/ERT2)Litt^*/WganJ, strain # 021160), ROSA26*^ZsGreen^* (B6.Cg-*Gt(ROSA)26Sor^tm6(CAG-ZsGreen1)Hze^*/J, strain # 007906), ROSA26*^tdTomato^* (B6.Cg-*Gt(ROSA)26Sor^tm9(CAG-tdTomato)Hze^*/J, strain # 007909), iDTR (C57BL/6-*Gt(ROSA)26Sor^tm1(HBEGF)Awai^*/J, strain # 007900), TRPV1-Cre (B6.129-*Trpv1^tm1(cre)Bbm^*/J, strain # 017769), *Trpv1^-/-^* (B6.129X1-*Trpv1^tm1Jul^*/J, strain # 003770), and NG2*^DsRed^* (Tg(Cspg4-DsRed.T1)1Akik/J, Strain # 008241). All transgenic mice were inbred in our laboratory, validated, and genotyped for the study. Both male and female mice were used. All animal experiments were approved by the Oregon Health & Science University Institutional Animal Care and Use Committee (IACUC IP00000968).

To generate CX3CR1^CreER^; ROSA26^ZsGreen^ mice, CX3CR1^CreER^ mice were crossed with ROSA26^ZsGreen^ mice. These mice were used to evaluate changes in the macrophage population following the perforation of the TM. CX3CR1^CreER^; ROSA26^tdTomato^ mice were created by breeding CX3CR1^CreER^ mice with ROSA26^tdTomato^ mice to assess the source of macrophage recruitment after TM perforation. Additionally, CX3CR1^CreER^; iDTR mice were developed by crossing CX3CR1^CreER^ mice with iDTR mice to facilitate macrophage depletion. To activate Cre-mediated recombination, tamoxifen (TAM) was administered via intraperitoneal (IP) injection at a dose of 75 mg/kg body weight every 24 hours for three consecutive days. For macrophage depletion, diphtheria toxin (DT) was administered intraperitoneally at a dose of 10 ng/kg body weight once every 24 hours for four consecutive days, starting one day after the administration of TM prior to TM perforation. iDTR mice served as control animals.

### TM perforation

Animals were anesthetized using an intraperitoneal injection of a ketamine (20 mg/mL) and xylazine (2 mg/mL) mixture in saline, at a dosage of 0.1 mL per 20 g of body weight. After achieving anesthesia, a sterile 26-gauge needle was employed to create a perforation in the pars tensa of the TM. After the animals awakened and fully recovered, they were returned to the animal facility. The procedure has been approved by the IACUC protocol under TR01_IP00000968.

### 5-Ethynyl-2’-deoxyuridine (EdU) labeling of TM

EdU (5 mg/mL in PBS, Sigma-Aldrich, Cat. # 900584) was administered intraperitoneally at a dose of 25mg/kg body weight starting one day prior to TM perforation and then once every two days until the animals were sacrificed (27). TMs were isolated and fixed in 4% PFA at 4°C overnight. The following day, proliferating cells in the TMs were assayed using a Click-iT® Plus EdU Alexa Fluor® 555 imaging kit (Life Technologies, Cat. # C10638) used according to the manufacturer’s instructions. Briefly, TMs were washed once in PBS, permeabilized in 1% Triton X-100 for 2 hours, and immunoblocked with 3% BSA in PBS for 30 minutes. Samples were then incubated in the EdU cocktail for 30 minutes, washed once in 3% BSA in PBS, mounted using mounting medium (Vector Laboratories, Cat. # H-1200) and visualized under an Olympus FV1000 laser-scanning confocal microscope (Olympus FV1000, Japan).

### Bone marrow transplantation

CX3CR1*^CreER^*; ROSA26*^tdTomato^*, C57/BL, and *TRPV1^-/-^* mice (6 weeks old, both sexes) were irradiated with a γ-emitting source at a dose of 9 Gy. Immediately following irradiation, the mice were reconstituted via a single periorbital sinus injection of 2 × 10^7^ fluorescent bone marrow derived cells (BMDCs) from donor CSF1R*^EGFP^* mice (4 weeks old) in 200 μl of modified Hank’s Balanced Salt Solution (HBSS). To prevent potential radiation effects on tissue-resident macrophages in the TM, the animals were protected with a specially designed head shield during irradiation (Mildner et al. 2007). After transplantation, the mice were provided with antibiotics in their drinking water (1.1g/L neomycin sulfateand 167mg/L polymyxin B sulfate) and monitored daily for four weeks. At four weeks post-transplantation, the recipient mice were used for further studies.

### Bulk RNA-seq analysis

Total RNA was extracted from the TM of wild-type (WT) mice, including both control and post-perforated groups, on days 1, 3, and 7. Three replicates were collected per group, with three TMs per sample, using the RNeasy Mini Kit (Qiagen, Cat. # 73404) following the manufacturer’s instructions. The concentration and quality of the isolated RNA samples were assessed using an Agilent TapeStation system. Only samples with RNA integrity numbers (RIN) greater than 8.5 were selected for library preparation and sequencing with polyA selection (40M/PE150). Data analysis was conducted using the ROSALIND® pipeline. Differentially expressed genes (DEGs) between the different groups (post-injury days 1, 3, and 7 compared to control) were evaluated. This evaluation included calculating fold changes, p-values, and optional covariate correction with DESeq2. Additionally, hypergeometric distribution analysis was performed to analyze gene ontology (GO) enrichment. The topGO R library was utilized to determine local similarities and dependencies between GO terms using an Elim pruning correction. Enrichment was calculated relative to a set of background genes relevant to the experiment.

### scRNA-seq analysis

Single-cell RNA sequencing data on murine TMs across several wound healing stages were obtained from the NCBI Gene Expression Omnibus (GEO; accession number GSE196692) (Scaria et al. 2023). The sequences were aligned to the mouse genome (mm10), and standard processing was conducted using Seurat v5 in R (Team 2020; Satija et al. 2015; Hao et al. 2024). Cells expressing fewer than 200 genes and genes expressed in fewer than three cells were excluded from the analysis. Data were log-normalized, and principal component analysis (PCA) was performed using the first 16 components for graph-based clustering, followed by Uniform Manifold Approximation and Projection (UMAP) for dimensionality reduction. Supervised classification was conducted using specific marker genes of macrophages and endothelial cells. Gene expression trends over time were visualized for selected genes. Additionally, the top 500 genes from each stage were analyzed using the PANTHER Classification System for PANTHER pathway and GO biological process enrichment (Mi et al. 2013; Thomas et al. 2003), revealing biological processes such as angiogenesis and wound healing. Macrophage-to-endothelial cell communication was analyzed using the CellChat package (Jin et al. 2021), identifying key signaling pathways. All visualizations were constructed to highlight differences in biological processes, gene expression patterns, and cell-cell communication across the wound healing timeline.

### Immunofluorescence in frozen sections of the TM and trigeminal ganglion

The auditory bulla and capsule, which enclose the TM, middle ear, and trigeminal ganglion, were isolated and fixed overnight in 4% paraformaldehyde (PFA). After washing three times with phosphate-buffered saline (PBS), the TMs were carefully dissected and incubated the following day in a permeabilization solution containing 1% Triton X-100 in PBS for 2 hours at room temperature (RT). Afterward, the tissues were blocked in a solution composed of 0.25% Triton X-100, 10% goat serum, and 1% bovine serum albumin (BSA) in PBS (blocking/permeabilization solution) for 1 hour at RT. The samples were then transferred to a primary antibody solution containing an anti-F4/80 monoclonal antibody (BM8) from eBioscience (Invitrogen Cat# 14-4801-85), diluted in the blocking/permeabilization solution, and incubated overnight at 4°C. The next day, after washing three times with the blocking/permeabilization solution, the samples were incubated for 1 hour at RT with a fluorescence-conjugated secondary antibody in the same blocking/permeabilization solution. Finally, the samples were washed three times with PBS, mounted on slides, and visualized under an Olympus laser-scanning confocal microscope (model number: FV1000, Japan).

The trigeminal ganglion was dehydrated sequentially in 15% and 30% sucrose solutions, frozen, and embedded in optimal cutting temperature (OCT) compound. Tissues were sectioned at a thickness of 10 μm for subsequent immunofluorescence staining. Specimens were incubated in blocking/permeabilization for 1 hour at RT, followed by incubation in primary antibody solution (diluted in blocking/permeabilization solution) overnight at 4°C. After three washes with PBS the next day, samples were incubated with the fluorescence-conjugated secondary antibody in the blocking/permeabilization solution for 1 hour at RT. Following three additional washes with PBS, samples were mounted using Antifade Mounting Medium with DAPI and visualized under an Olympus FV1000 laser-scanning confocal microscope (Olympus FV1000, Japan).

### Macrophage quantification

The TM was imaged using an Olympus FV1000 laser-scanning confocal microscope (Olympus FV1000, Japan) equipped with a 10x objective lens. Composite images were created by stitching together four tiles to capture the entire TM. To ensure consistent comparability across all confocal scans, laser intensity, step size, and resolution were maintained throughout the imaging process. Macrophage numbers in the wound area were quantified using the 3D object counter in Fiji software (ImageJ, NIH, version 1.51t). A thresholding technique was applied to distinguish object voxels from background voxels. Additionally, a size filter was set between 300 to 1500 μm³ to exclude objects outside this defined range from the analysis. The density of tissue-resident macrophages (cells per mm²) was calculated by dividing the total number of macrophages by the area of the TM analyzed. Mean values were determined for each sample and subsequently analyzed and compared across different groups.

### Assessment of vascular density

Lectin-DyLight 649 (Vector Laboratories, Cat. # DL-1178-1) was diluted in 0.1 M PBS buffer to a concentration of 0.5 mg/ml (vol. 100 µl) and administered to anesthetized mice via intravenous retro-orbital sinus 10 minutes prior to sacrifice. TMs were isolated and fixed overnight at 4°C in 4% PFA. The following day, TMs were mounted and visualized under an Olympus FV1000confocal microscope with a 10x objective lens. The blood vessel and TM areas, vessel diameters, and fluorescence intensities in the vessels and perivascular interstitium were quantified using Fiji. software. Vascular density was defined as 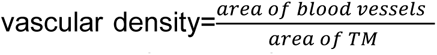. Vascular permeability was calculated as 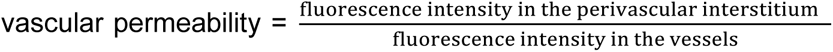 (Shi et al. 2014). The vascular diameters and permeability of three vessels surrounding the wound area were measured in each sample.

### qRT-PCR

Total RNA samples were reverse-transcribed using a RETROscript kit (Ambion), and cDNA was synthesized from total RNA diluted 10-fold in DNase-free water. Each cDNA sample was independently measured three times. Transcripts were quantified by TaqMan Gene Expression Assay for *Tac1* (Mm01166996_m1) on an Applied Biosystems 7300 real-time PCR system. The real-time PCR was cycled at 95°C for 20s, followed by 40 cycles at 95°C for 1s and 60°C for 20s. Mouse GAPDH served as the endogenous control. Quantitative analysis was performed according to the guidelines provided by Applied Biosystems using the comparative cycle threshold method.

### Statistics

All experiments were designed with proper controls. Statistical analyses were performed using GraphPad Prism 9 software (GraphPad Software). Statistical difference between two groups was evaluated by unpaired, two-tailed t test. One-way ANOVA followed by Tukey’s multiple comparison test was used to compare differences across multiple groups. Differences were considered significant at p<0.05. Data were presented as the mean ± SEM.

## Results

### Macrophages ‘gathering’ in the vicinity of the injured TM wound region are primarily recruited from blood-circulating monocytes and not locally proliferated or migrated

In this study, a perforation was created in the antero-inferior quadrant of the pars tensa of the TM in an adult CX3CR1^CreER^; ROSA26^ZsGreen^ mouse model. The antero-inferior quadrant was chosen because it is the most common site for traumatic TM perforations seen in clinic. In this model, CreER-mediated recombination was activated by tamoxifen injections, enabling visualization of Cx3cr1+ cells through green GFP fluorescence. Following the TM perforation, we observed that macrophages began to migrate toward the perforation site by day 3. By day 7, the injured membrane was largely covered by an accumulation of macrophages (Figure 1A and B). This raises the question: What is the source of these macrophages? We propose three possible sources that could explain the increased population of macrophages: recruited macrophages from non-wounded areas, migrating monocytes from the bloodstream, or proliferating local tissue-resident macrophages. To determine the source, we first administered EdU, a nucleoside analog, to the animals one day before perforation and then once every two days after the perforation until the point of sacrifice. We observed very little overlap between the ZsGreen+ and EdU+ signals, as shown in the higher magnification images (Figure 1A, lower panel). This finding indicated that the accumulated macrophages were recruited from distant sources rather than locally proliferated. Next, we developed a chimeric bone marrow transplantation (BMT) and fluorescence reporter mouse model (Figure 1C). In this model, CSF1R^EGFP^ donor bone marrow cells were transplanted into CX3CR1^CreER^; ROSA^26tdTomato^ recipient mice. This allowed us to distinguish local tissue-resident macrophages (tdTomato+) from blood-circulating monocytes (EGFP+). One month after transplantation, when the mice were effectively reconstituted (Park et al. 2021), we perforated the TMs of the recipient mice. After perforation, we observed a significant presence of EGFP+ cells recruited to the wound area, while few tdTomato+ cells were found (Figure 1D). This experiment further confirmed that the macrophages accumulating around the wound primarily originated from blood-circulating monocytes, rather than from locally proliferated or migrated tissue-resident macrophages.

**Figure 1.**
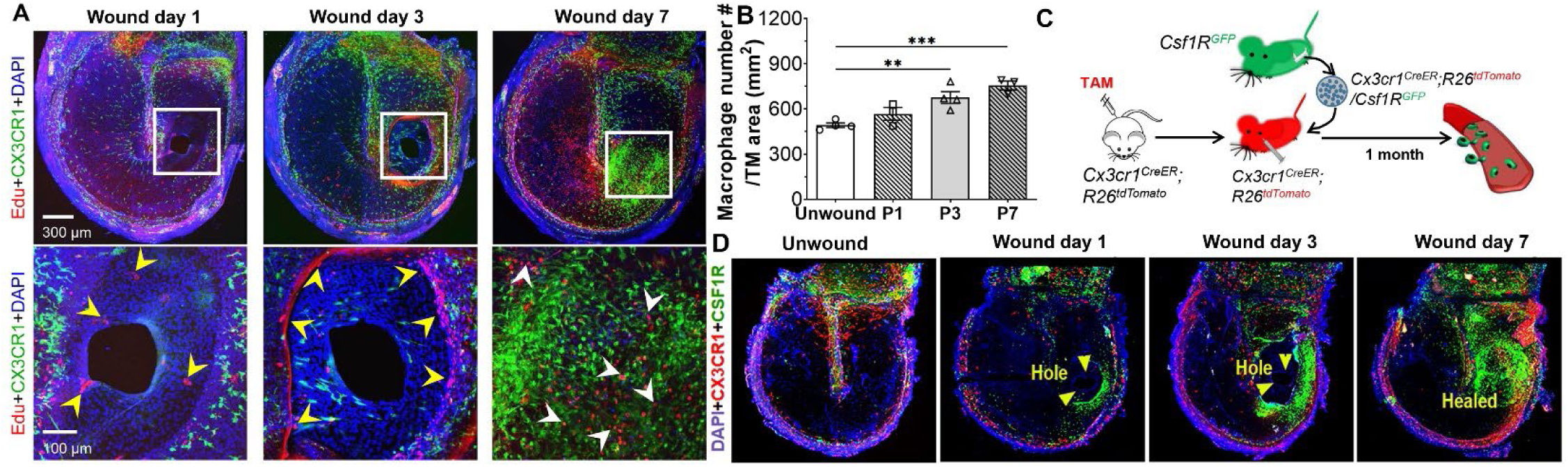
Blood-derived monocytes, not local tissue-resident macrophages, predominate in TM wound sites. (**A**) The perforated TM was healed within a week, as shown in the low magnification (upper, right) and high magnification (lower, right) confocal images. EdU pause labeling of the proliferating cells (arrowheads) *in vivo* shows some, but only a few of the macrophages, incorporate EdU (arrowheads in lower right panel, EdU+ cells). (**B**) Statistical analysis shows that a significant population of macrophages ‘gather’ in the injured areas of the perforated TM during wound healing as shown on day 3 (676 ± 39 cells/mm^2^, n = 4) and day 7 (756 ± 29 cells/mm^2^, n = 3), compared to non-perforated TMs (492 ± 15 cells/mm^2^, n = 4) and on day 1 (567 ± 42 cells/mm^2^, n = 3). One-way ANOVA followed by Tukey’s multiple comparison test, **p<0.01, ***p<0.001. Data are presented as a mean ± SEM. (**C**) Schematic representations illustrate how the chimeric mouse model was created. (**D**) Confocal images of TMs collected from CSR1F^EGFP^ bone marrow transplanted CX3CR1^CreER^; ROSA26^tdTomato^ mice before and on days 1, 3, and 7 after perforation. Blood-derived monocytes (tagged by EGFP under a Csf1r promoter (green), not local tissue-resident macrophages (red), show marked accumulation in the wound sites.

### Angiogenesis takes place during wound healing

Angiogenesis is crucial for wound healing, and numerous studies have highlighted the link between impaired angiogenesis and delayed wound recovery (Schugart et al. 2008). In our study, we observed early vascular inflammation (Figure 1, Supplementary Materials) and subsequent angiogenesis occurring alongside the recruitment of monocytes near the wound site. Angiogenesis began on day 3 after perforation and continued to progress by day 7 (Figure 2A-C). We noted a significant increase in vascular density around the wound area (Figure 2E). The newly formed immature vessels were large in diameter and showed increased permeability, as evidenced by the extravascular presence of circulating fluorescently labeled lectin (Figure 2F and G). Using a fluorescent reporter mouse model, we found that pericytes (NG2DsRed) migrated and proliferated as tip cells during angiogenesis (Figure 2D, right panels a, b). This finding aligns with our previous report on NG2+ pericyte-led neovascularization in the inner ear (Zhang et al. 2021). These observations suggest that sprouting angiogenesis, mediated by pericytes, occurred during TM wound healing.

**Figure 2.**
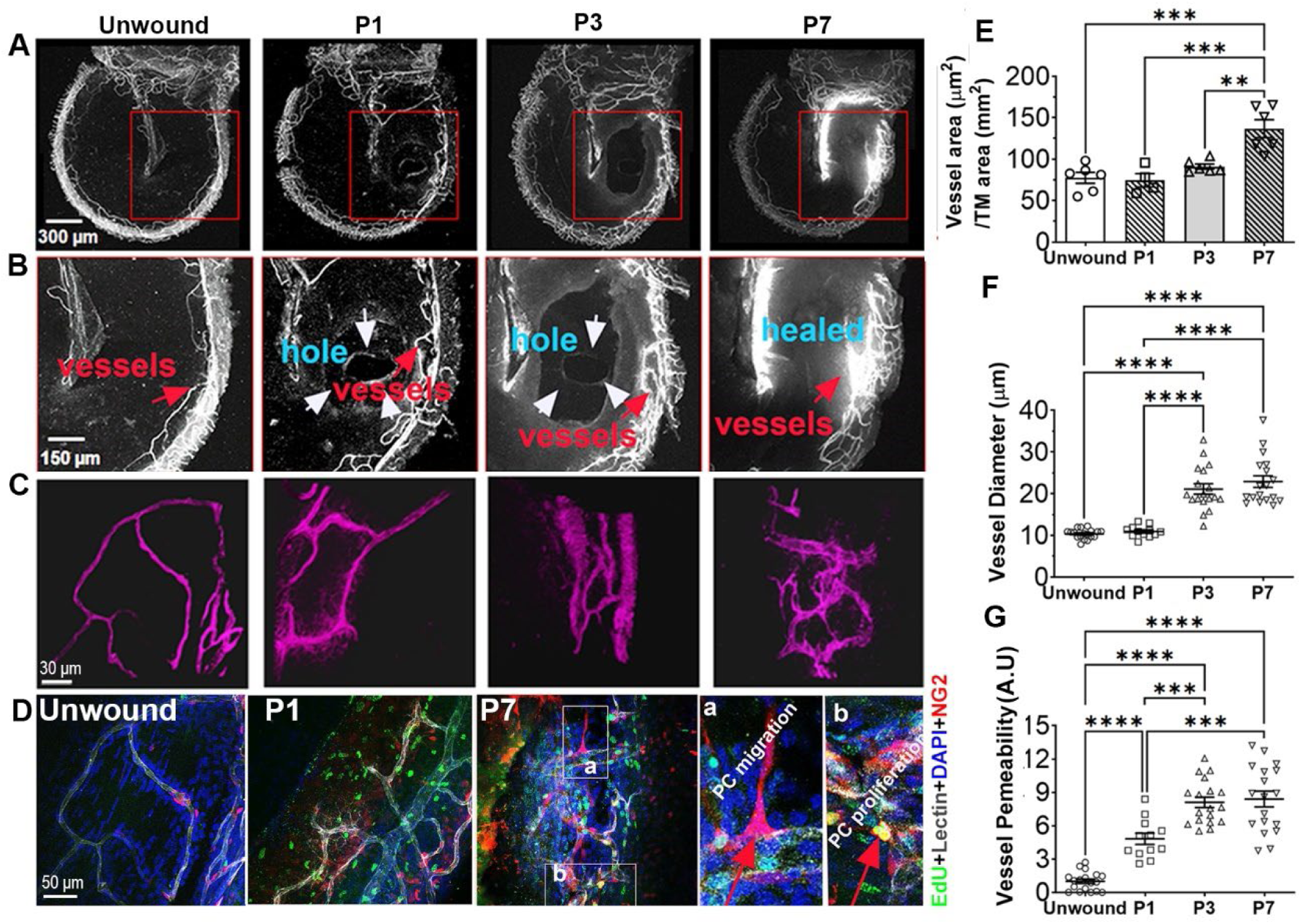
Angiogenesis in the vicinity of vessels in the perforated TM. (**A** and **B**) Angiogenesis is seen near wound areas by day 3 and progressed by day 7 after TM injury, shown under low (**A**) and high magnification (**B**). (C) 3D reconstructions of the vascular structure on different days after perforation. (**D**) No EdU+ signal was seen in vascular cells of the TMs in the NG2^DsRed^ mouse (Control). In contrast, increased EdU+ cells (green), including vascular cells, were seen on day 1 (P1) and day 7 (P7) following perforation. Pericyte migration (to form tip cells) and proliferation from vessels are better visualized in the zoomed images (a and b). (**E**) Statistical analysis of vessel density in the wound region of normal (n = 6) and perforated TMs on day 1 (n = 4), day 3 (n = 6), and day 7 (n = 6). (**F**) Statistical analysis of vessel diameter in the wound region of normal (n = 18) and perforated TMs on day 1 (n = 12), day 3 (n = 18), and day 7 (n = 18). (**G**) Statistical analysis of vessel permeability in the wound region of perforated TMs on day 1 (n = 12), day 3 (8.1 ± 0.47 μm, n = 18), and day 7 (8.41 ± 0.71 μm, n = 18). One-way ANOVA followed by Tukey’s multiple comparison test, *p<0.05, **p<0.01, ***p<0.001, ****p<0.0001. Data are presented as a mean ± SEM.

### Macrophages promote angiogenesis and depletion of macrophages attenuates angiogenesis

Monocyte recruitment and angiogenesis occur simultaneously during wound healing in the TM, as we have demonstrated. Study has shown that macrophages play a key role in stimulating vessel sprouting during wound angiogenesis in zebrafish (Gurevich et al. 2018). To investigate the role of macrophages in angiogenesis in adult mice, we developed an inducible macrophage depletion model (CX3CR1CreER; iDTR). This model was created by crossing CX3CR1^CreER^ transgenic mice with an inducible diphtheria toxin receptor (iDTR) mouse line that carries a Cre-dependent simian diphtheria toxin receptor. This combination enables targeted cell death upon the administration of diphtheria toxin (Zhang et al. 2021; Buch et al. 2005). At one month of age, the mice were treated with tamoxifen (TAM) for three days to induce the expression of Cre recombinase. To deplete macrophages, DT was administered through daily intraperitoneal injections at a dose of 10 µg/kg for four consecutive days, starting one day after TAM treatment (Zhang et al. 2021), followed by an acute perforation of the TM in the pars tensa. Our results showed that, compared to control mice (Figure 3A), the recruitment and accumulation of macrophages (labeled with F4/80) in the wound area were significantly reduced at day 3 post-perforation in the macrophage depletion mice (Figure 3B). By day 7, the wound area in mice that were not depleted of macrophages was filled with macrophages and newly formed blood vessels (Figure 3C). In contrast, the macrophage-depleted mice showed an unhealed wound with reduced macrophage infiltration and less angiogenesis (Figure 3D). Quantitative analysis confirmed that both the macrophage population and vascular density were significantly lower in the macrophage-depleted mice at both day 3 and day 7 following perforation. Our data indicate that the depletion of macrophages leads to decreased angiogenesis and delayed wound closure during TM wound healing.

**Figure 3.**
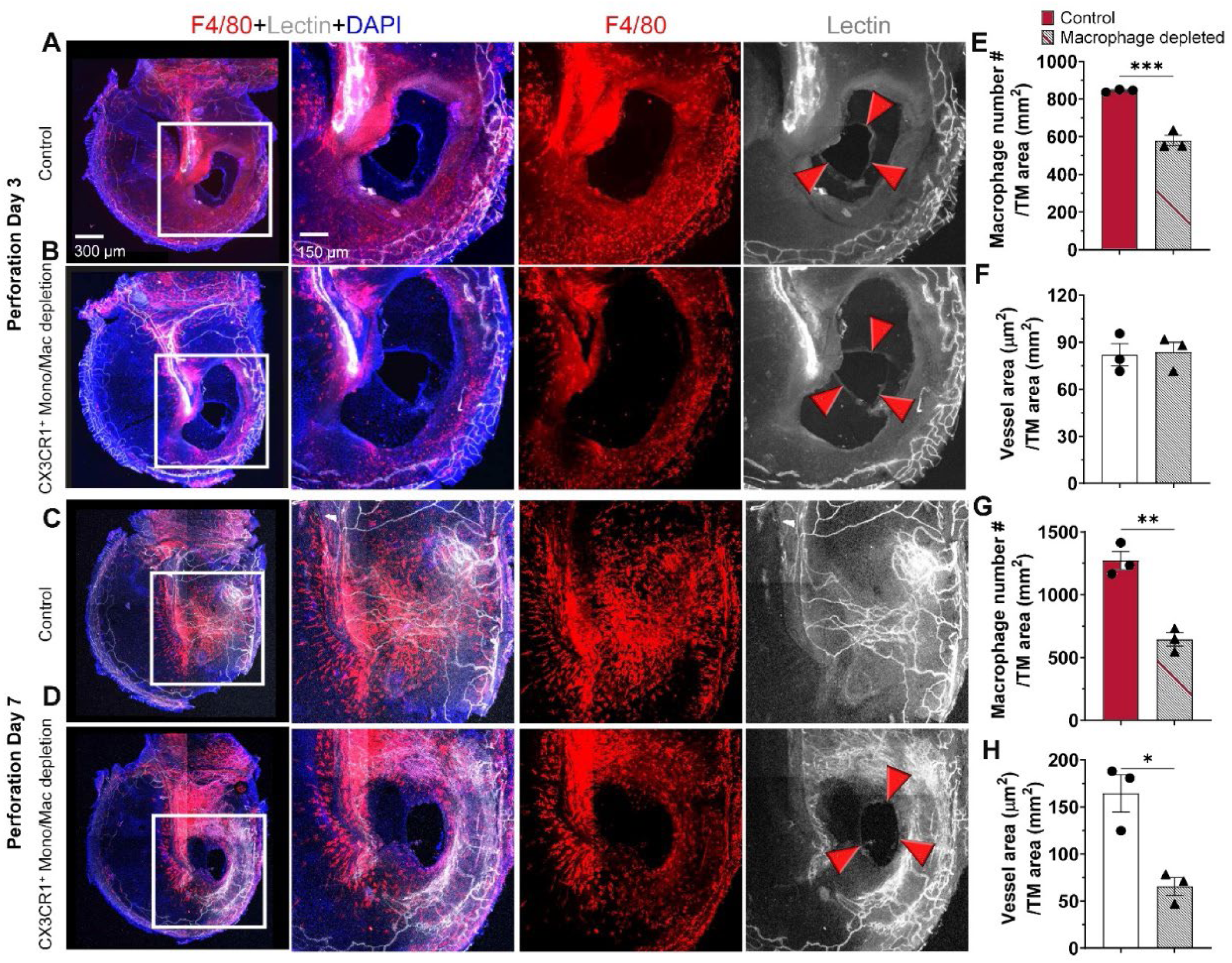
Depletion of macrophages attenuates angiogenesis and wound healing in the TM. (**A-D**) The representative images show macrophage accumulation and angiogenesis in the TM wound area, observed under both low (left panel) and high magnifications (middle and right panels) in non-macrophage-depleted (control) and macrophage-depleted mice at 3- and 7-days following perforation. (**E-H**) Quantitative analysis indicates a significant reduction in both the macrophage population and vascular density in the macrophage-depleted mice by day 3 and day 7 post-perforation (n = 3, unpaired t-test). *p<0.05, **p<0.01, ***p<0.001. Data are presented as mean ± SEM.

### Angiogenesis is linked to inflammation, and multiple molecular signals are associated with macrophage-driven angiogenesis

What are molecular signals are associated with macrophages-driven angiogenesis? Angiogenesis in the provisional matrix during wound healing is affected by various factors, including tissue hypoxia and the cytokines produced by macrophages (Veith et al. 2019). With bulk RNA-seq analysis, on a large picture, we also identified the parallel enrichment of Gene Ontology (GO) biological processes, including inflammatory response (GO: 0006954), leukocyte migration involved in inflammatory response (GO: 0002523), regulation of angiogenesis (GO: 0045765), angiogenesis (GO: 0001525), and sprouting angiogenesis (GO: 0002040) at different time points after perforation of the TM (Figure 4A-C). Figure 4D-J specifically show differential gene expression (DEGs) involved in the angiogenetic signaling pathways at different times during TM wound healing, which include R*dpjl, Hif1a, Pla2g4d, Vegfa, Pik3r3, and Rhoc.* molecular signals.

**Figure 4.**
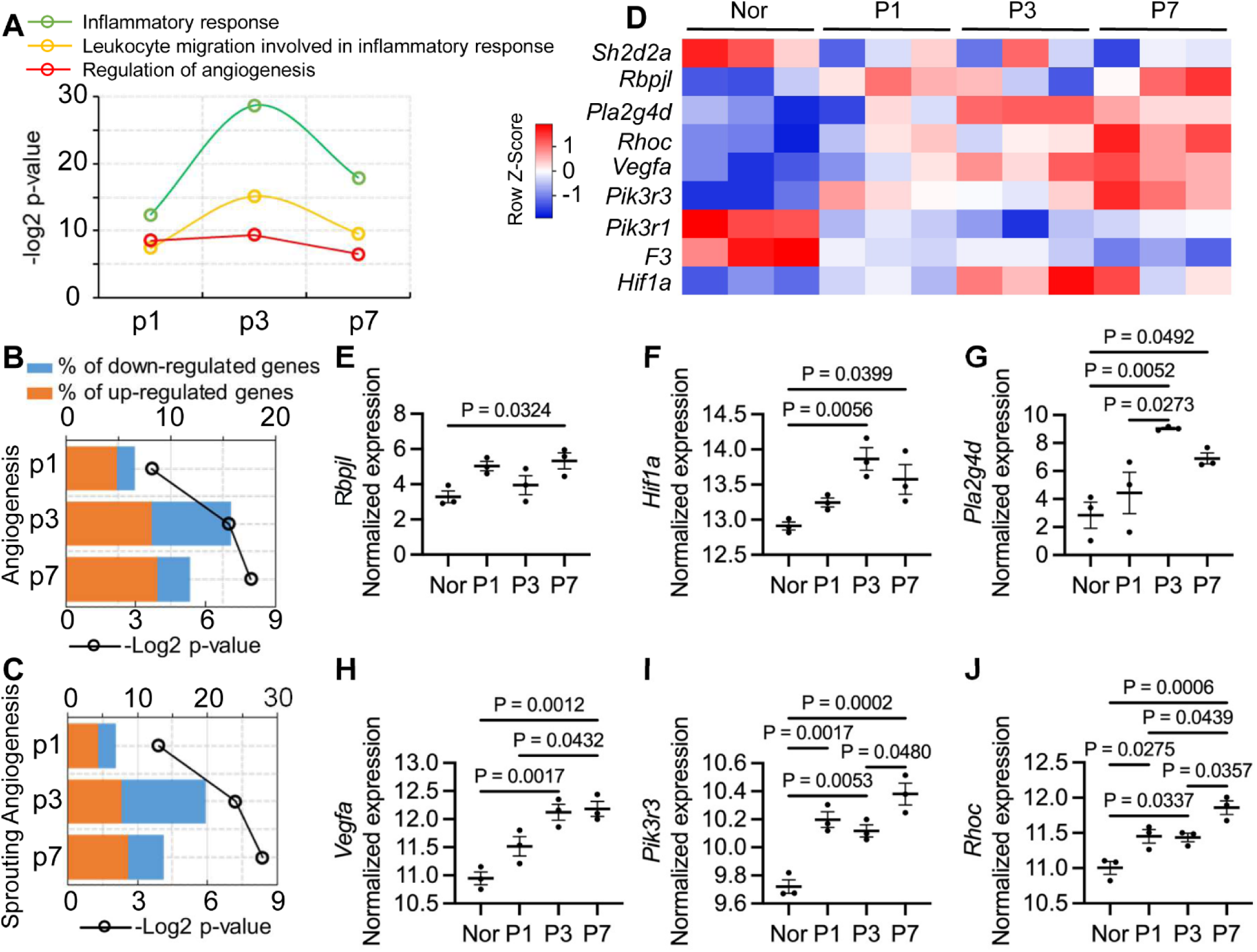
Gene ontology enrichment analysis and heat map of DEGs in normal and perforated groups at different time points. (**A-C**) ROSALIND® analysis shows the hypergeometric distribution of the enrichment of GO biological processes including inflammatory response, leukocyte migration involved in inflammatory response, regulation of angiogenesis, angiogenesis, and sprouting angiogenesis at different time points after TM perforation. (**D-J**) Heat map analysis (**D**) and representatives (**E-J**) of individual DGEs (relative to the angiogenesis signaling pathway) in control and perforated groups at different time points shows a significant increase by days 3 and 7 after perforation (n = 3). One-way ANOVA followed by Tukey’s multiple comparison test. Data are presented as a mean ± SEM.

To further investigate the roles of macrophages in driving sprouting angiogenesis, we re-analyzed a published sc-RNA-seq dataset (GEO dataset: GSE196692) collected from mouse TMs at various time points following TM perforation (Scaria et al. 2023). Our focus was primarily on the signature genes of macrophage clusters. Notably, the gene ontology enrichment analysis of these macrophage clusters revealed significant enrichment of genes associated with angiogenesis, epidermal development, keratinization, and extracellular matrix organization at different stages of TM wound healing (Figure 5B-D). The role of macrophages in angiogenesis is particularly highlighted in Figure 5E. Furthermore, we identified macrophage involvement in angiogenic activity linked with the *Hif1a, Vegfa, Pla2g4d, Rdpjl, Pik3r3*, and *Rhoc* pathways (Figure 5F). These signals correspond closely to the pathways identified through bulk gene analysis as shown in Figure 4E-J. By integrating single-cell RNA sequencing data with spatial information, we observed active interactions between macrophages and endothelial cells during TM wound healing (Figure 5G and H). Notably, from post-injury day 1 to day 7, there was significant intercellular communication between macrophages and endothelial cells mediated by secreted phosphoprotein 1 (*Spp1*), a signaling molecule that promotes angiogenesis in other organ systems (Tsai et al. 2020; Rowe et al. 2014). In contrast, no interactions between macrophages and endothelial cells were observed in unwounded tissues (Figure 5G). Taken together, our data indicate angiogenesis is mediated by multiple molecular signals and closely associated with inflammation and macrophages during wound healing.

**Figure 5.**
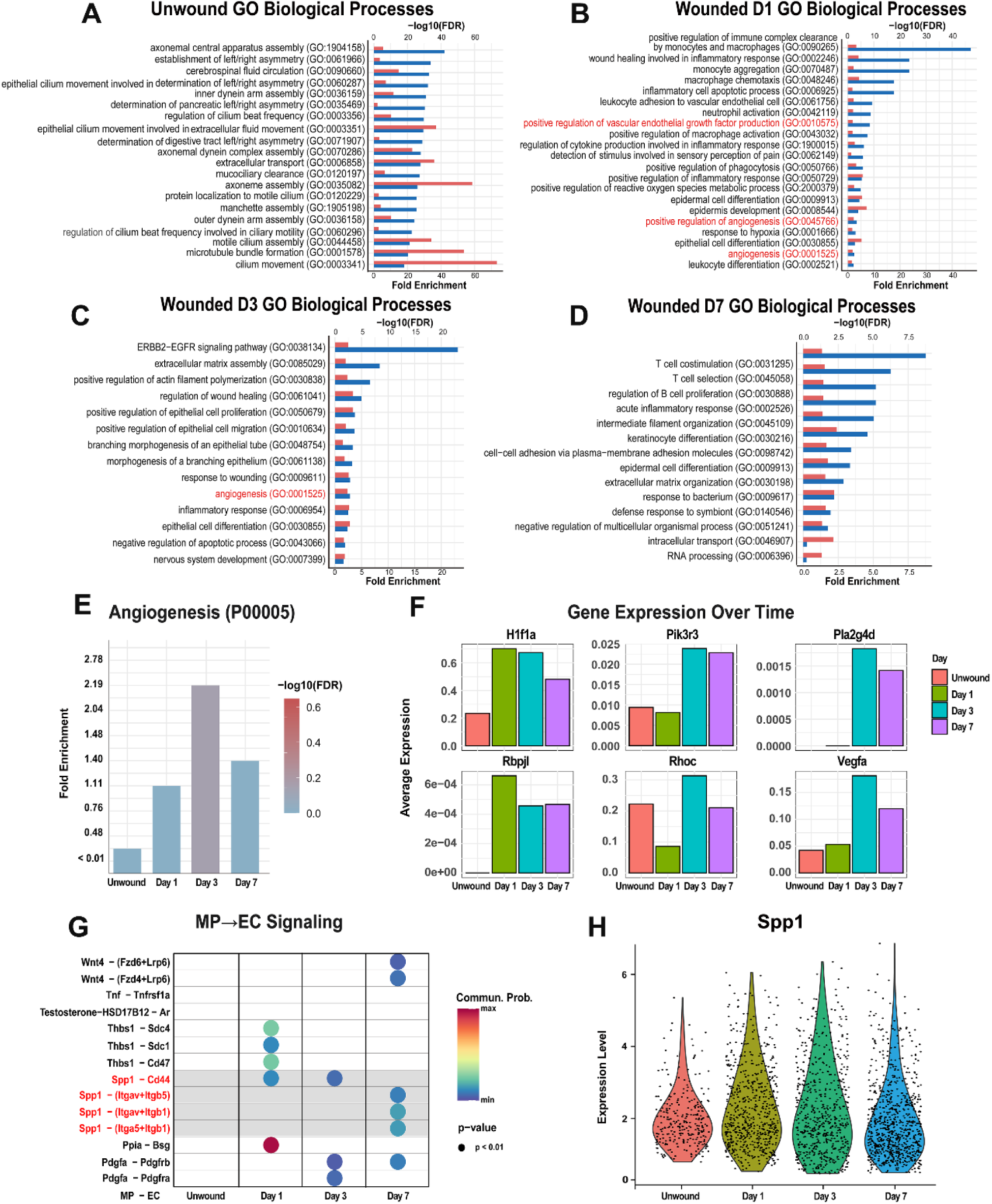
Sc RNA sequencing analysis shows dynamics of gene ontology, pathways, gene expression and cell-cell interactions during wound healing in the murine TM. (**A-D**) GO enrichment analysis across different time points (**A**-Unwound, **B**-Day 1, **C**-Day 3, and **D**-Day 7), with blue bars representing fold enrichment and red bars representing statistical significance based on −log10(FDR), highlighting key processes of macrophages involved in wound healing. (**E**) PANTHER Pathway analysis of macrophage signature genes shows the enrichment of Angiogenesis (P00005). (**F**) Average expression of selected macrophage genes (*Hif1a, Pik3r3, Pla2g4d, Rhoc, Rbpj, and Vegfa*) over time, demonstrating their dynamic regulation during wound healing. (**G**) Cell-cell communication networks between macrophage (MP) and endothelial cell (EC) reveal MP→EC signaling mediated by *Spp1* across wound healing stages. (**H**) Violin plots show the expression levels of *Spp1* in macrophages at each wound healing stage.

### Monocyte recruitment in wound TM is mediated by TRPV1 signal

The final central question we asked in this study was: what molecular signals trigger monocyte migration to the wound area during TM wound healing? The recruitment of macrophages is known to be complex and tissue specific. Recent studies have emphasized the role of interactions between neurons and the immune system in mediating wound healing processes (Wang et al. 2020; Lowy et al. 2021). Through bulk RNA sequencing analysis of normal and perforated TMs at various time points, we identified an enrichment of the neuroinflammation response process (Gene Ontology: 0150076) in perforated TMs (Figure 6 A and B). Notably, we observed a significant upregulation of the Tac1 gene—up to tenfold compared to non-perforated conditions (Figure 6C). The Tac1 gene encodes several neuropeptides from the tachykinin family, including substance P, neurokinin A, neuropeptide K, and neuropeptide gamma, all of which are closely linked to TRPV1 signaling (Devesa et al. 2014). The TM is a highly sensitive tissue rich in sensory nerve fibers. A rupture of the TM can result in intense pain for patients. The TRPV1 channel is well-known for its crucial role in pain sensation (Marrone et al. 2017). TRPV1-dependent pain signals can rapidly trigger vascular inflammation, immune responses, and the proliferation of precursor cells. Using a fluorescent reporter mouse line for TRPV1, we were surprised to discover that the TM is abundant in TRPV1-positive nerve fibers (Figure 6D). We further validated this finding in the trigeminal ganglion tissue (Figure 6E)(Wietecha et al. 2013).

**Figure 6.**
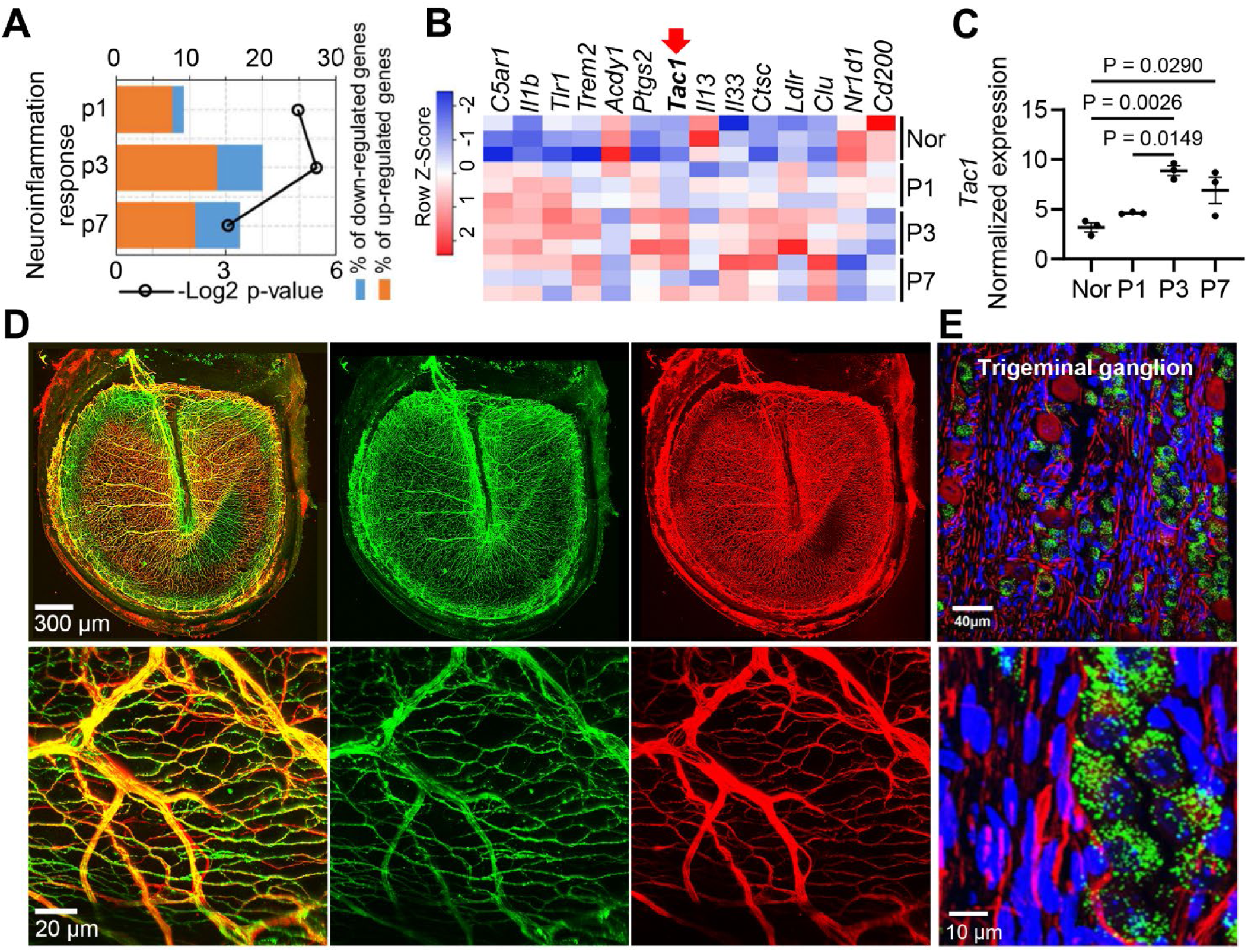
Heat map and GO enrichment analysis of DEGs revealed neuroinflammation response following TM injury and TRPV1 is richly expressed in the TMs of transgenic fluorescence TRPV1 reporter mice (*Trpv1^Cre^; R26^ZsGreen1^*). (**A**) ROSALIND® analysis shows hypergeometric distribution of the enrichment of the neuroinflammatory response at different time points after perforation of the TM. (**B**) Heat map of DEGs related to neuroinflammation in normal and perforated groups at different time points. (**C**) Bulk RNA-seq analysis shows a significant increase in Tac1 expression by days 3 and day 7 after perforation (n = 3). One-way ANOVA followed by Tukey’s multiple comparison test. Data are presented as a mean ± SEM. (**D** and **E**) TRPV1 expression in the TM and trigeminal ganglion under low (upper panels) and high magnification (low panels) in a TRPV1 fluorescence reporter mouse line (Trpv1^Cre^; R26^ZsGreen1^).

To evaluate the role of TRPV1 signaling in monocyte recruitment and angiogenesis, we examined TM wound healing in WT and *Trpv1^-/-^* mice transplanted with CSF1R^EGFP^ donor bone marrow cells. We observed that both monocyte recruitment and angiogenesis were markedly and significantly attenuated in *Trpv1^-/-^* mice on day 7 (Figure 7A-I). Perforated TM in *Trpv1^-/-^* mouse was only partially healed by day 7 and displayed reduced CSF1R^EGFP^ macrophages in the wound area (Figure7D and E). In contrast, WT mice exhibited robust monocyte recruitment (Figure 7C) and angiogenetic activity (Figure 7F). Perforated TM was healed by day 7 (Figure 7C). Additionally, we found that *Tac1* expression was significantly lower in *Trpv1^-/-^* mice by day 3 after perforation (see Figure 7J). These findings indicate a sensory neuronal-inflammatory interaction in TM wound healing, analogous to mechanisms observed in skin tissue (Zeng et al. 2024).

**Figure 7.**
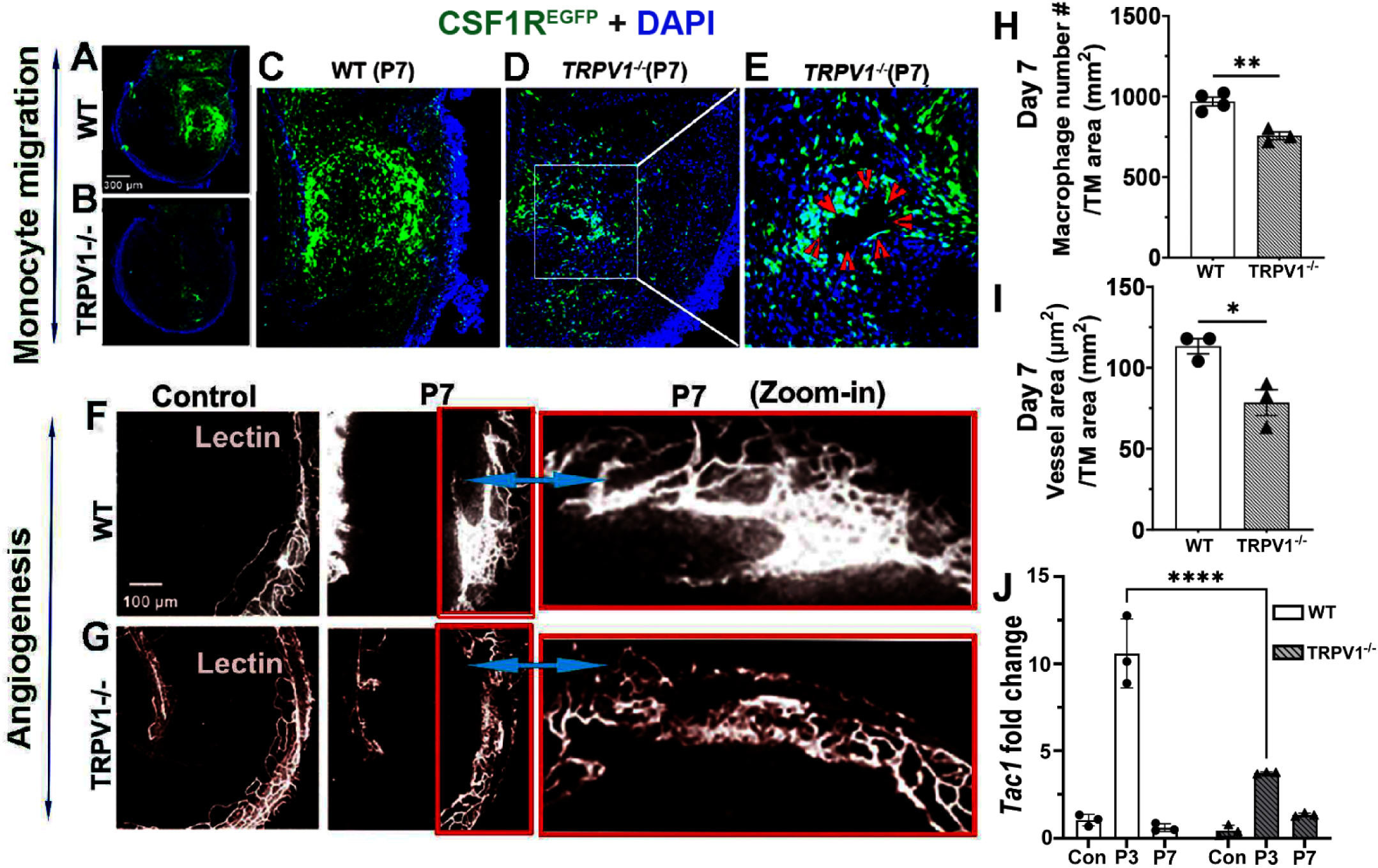
Reduced monocyte recruitment and angiogenetic activity in the perforated TM of chimeric Trpv1^-/-^ mice relative to the chimeric WT. (**A**) Perforated TM in the WT is fully healed on day 7, and the hole is massively covered with blood-derived CSF1R^EGFP^ macrophages. (**B**) Perforated TM in the *Trpv1^-/-^* mouse was not healed by day 7 and displayed reduced CSF1R^EGFP^ macrophages in the wound area. (**C** and **D**) High magnification images better show the difference in the level of monocyte migration in the WT and *Trpv1^-/-^* mice. (**E)** The Zoomed image from selected area in Panel D further shows the remaining unhealed perforation on day 7 (see red arrowheads). (**F**) and (**G**) (**F**, **G**, left panels) shows normal blood vessel distribution in the TM of WT and *Trpv1^-/-^* mouse TM before injury. (**F**, Middle panel) shows noticeable angiogenesis around wound areas in WT mice on day 7 after TM injury. (**G**, middle panel) shows that no apparent angiogenesis in the *Trpv1^-/-^* mice by day 7. (**F**, **G**, right panels) are high magnification images from selected areas of middle panels better show the difference between WT and *Trpv1^-/-^* animals. (**H**) There is a significant difference in the monocyte population in the wound TMs of the WT and *Trpv1^-/-^* mice (unpaired t-test, n = 3, **p<0.01). (**I**) There is also a significant difference in vessel density on day 7 following TM perforation between the WT and *Trpv1^-/-^* mice (n = 3, unpaired t-test, *p<0.05). (**J**) *Tac1* expression was significantly diminished in *Trpv1^-/^*^-^ mice by day 3 post-perforation (n = 3, p<0.0001).

## Discussion

This study demonstrates that acute TM wound healing involves macrophage recruitment and angiogenesis mediated by TRPV1 signaling. Specifically, we observed that macrophages derived from blood-circulating monocytes are rapidly drawn to the wound area following perforation. In parallel with the macrophage recruitment, we captured the dynamics of angiogenesis including early vascular inflammation and later vascular sprouting near the wound. Using bulk RNA gene analysis, we identified the DEGs enriched in the gene ontology at different time points after TM perforation, particularly regarding neuroinflammation response and angiogenesis. Further, by reprocessing published scRNA-seq datasets on unwounded and wounded murine TMs, we obtained the picture of vital biological processes that macrophages participate in during TM wound healing, including intercellular signaling between macrophages and endothelial cells and macrophage-associated angiogenesis. Finally, we found that the natural process of eardrum wound healing requires TRPV1-associated neuronal signaling, as *Trpv1-/-* mice display delayed or impaired wound healing with reduced monocyte recruitment and diminished angiogenic activity.

### Monocyte-derived macrophage recruitment and angiogenesis in TM wound healing

Macrophages are critical in tissue development, homeostasis, and repair of injury (Mosser et al. 2021). These innate immune cells are among the first responders to tissue injury and are vital at every stage of wound healing. They play a critical role in all phases of wound healing, from initiating inflammation to combat pathogen invasion, to resolving inflammation once the pathogens are eliminated, and further guiding vascular remodeling and stimulating local stem or progenitor cells to promote tissue remodeling and regeneration (Kim and Nair 2019; Gurevich et al. 2018; Pang et al. 2022; Sim et al. 2022). In our study, we observed a significant accumulation of macrophages at the site of the TM wound. During wound healing, macrophages can originate from different sources. They can rise from monocyte infiltration, proliferate locally, or migrate from other unwound areas in response to early tissue inflammation (Guilliams et al. 2013; Davies et al. 2013; Ginhoux and Guilliams 2016). Our findings indicate that the macrophages in wound tissue are primarily derived from blood-circulating monocytes, rather than from proliferating tissue-resident macrophages or significantly migrating from other tissue regions as shown in Fig.1. This observation aligns with previous studies that have identified the role of monocyte-derived macrophages in wound healing and tissue repair in various other tissues (Willenborg et al. 2012; Little et al. 2014). Research indicates that during the healing process, infiltrated monocytes first differentiate into M1 macrophages. These M1 macrophages play a crucial role in scavenging, phagocytosis, and antigen presentation during the inflammatory phase (Raziyeva et al. 2021). Over time, the gene expression of macrophages gradually shifts from the M1 phenotype to the M2 phenotype. This transition enables them to recruit stem cells and secrete cytokines and growth factors that stimulate the proliferation, differentiation, and migration of keratinocytes, fibroblasts, and endothelial cells (Kuraitis et al. 2022). In our study, we did not examine the plasticity of macrophage phenotypes during wound healing, nor did we investigate the specific roles of monocyte-derived macrophages throughout all phases of the healing process. Future studies that focus on identifying macrophage phenotypes in TM wound healing would be valuable. Such research could help unlock the therapeutic potential of manipulating specific macrophage functions to enhance TM wound healing.

Alongside macrophage recruitment, we observed early vascular inflammation (Figure 1, Supplementary Information) and later angiogenesis near the wound. Angiogenesis was initiated by day 3 post-perforation, and further progressed by day 7. Both macrophage recruitment and angiogenesis are critical for the early phase of the wound healing (Pang et al. 2022; Sim et al. 2022; Veith et al. 2019). Angiogenesis is particularly vital as it restores the supply of nutrients and oxygen to the wound area, aids in waste removal, and facilitates the transport of inflammatory cells, including monocytes (Gurevich et al. 2018). Impaired angiogenesis could result in stunted formation of granulation tissue and prolonged wound healing (Brem and Tomic-Canic 2007; Nunan et al. 2014). In our findings, we demonstrated that depleting macrophages resulted in reduced angiogenesis and delayed wound closure, as illustrated in Figure 3. Accumulated evidence indicates that macrophages secrete various pro- and anti-angiogenic growth factors, which are crucial for stimulating angiogenesis during tissue repair and wound healing (Corliss et al. 2016; Stockmann et al. 2011).

Angiogenesis is a complex process that involves various cell types and a coordinated sequence of events within the microenvironment, including vascular sprouting, regression, and maturation (Shi et al. 2023; Wietecha et al. 2013). Endothelial cells play a pivotal role at the cellular level, serving as the foundational scaffold during tissue repair. As angiogenic sprouting progresses, these cells undergo processes such as activation, adhesion, proliferation, and migration (Wacker and Gerhardt 2011; Shi et al. 2023). In our previous study on the inner ear, we found that pericytes act as tip cells and are essential for guiding vessel sprouting (Zhang et al. 2021). Consistent with these prior findings, we observed that angiogenesis in the wounded TM is driven by the migration and proliferation of NG2+ pericytes. At the molecular level, wound angiogenesis is regulated by signals from both serum and the surrounding extracellular matrix (ECM) (Li et al. 2003). Several angiogenic growth factors have been shown to accelerate healing of TM wounds (Uddman et al. 1988).

A previous study demonstrated that the topical application of epidermal growth factor (EGF) effectively heals chronic TM perforations in an animal model (Bai et al. 2023). In our bulk gene analysis, we observed significant upregulation of several genes associated with angiogenesis, including *Rdpjl, Hif1a, Pla2g4d, Vegfa, Pik3r3,* and *Rhoc*. While we did not thoroughly investigate the specific molecular signals related to macrophage-mediated angiogenesis in the wounded TM in this study, we reanalyzed a previously published scRNA-seq dataset (GEO dataset: GSE196692) (Scaria et al. 2023). This dataset was collected from mouse TMs at various time points following TM perforation (Scaria et al. 2023). Our primary focus was on the signature genes from macrophage clusters. Gene enrichment analysis of these macrophage clusters revealed a high enrichment of transcripts related to angiogenesis, epidermal development, keratinization, and extracellular matrix organization (Figure 5A-D). The macrophage-related angiogenesis was associated with *Rdpjl, Hif1a, Pla2g4d, Vegfa, Pik3r3*, and *Rhoc* signals. These signals were consistently identified in our bulk gene analysis of the entire TM preparation. Most interestingly, from post-perforation days 1 to 7, we observed strong intercellular communication between macrophages and endothelial cells through phosphoprotein 1 (SPP1). Notably, no SPP1-mediated macrophage-to-endothelial cell interactions were detected in the unwounded tissues (Figure 5G).

SPP1, also known as osteopontin, is a secreted multifunctional glyco-phosphoprotein that is upregulated in acute and chronic inflammatory settings, signaling via integrin and CD44 receptors (Yim et al. 2022). An early study demonstrated that microglia are a major source of SPP1 under stress conditions, contributing to angiogenesis (Waller et al. 2010). Other studies have shown that SPP1 contributes to angiogenesis by upregulating VEGF signaling pathways (Takano et al. 2000; Lou et al. 2021). However, the current underlying regulatory mechanisms of *Spp1* signaling in macrophage-EC-associated angiogenesis remain poorly understood. In this study, we did not examine the individual effects of specific signals on angiogenesis, nor did we investigate whether blocking the SPP1 signal would impact the identified angiogenic factors. It is clear that future research is needed to explore the role of SPP1+ macrophage-associated angiogenesis in wound TM repair. This research would be highly clinically relevant, as it could enhance blood circulation and improve wound healing.

### TRPV1 mediated neuronal inflammation facilitates TM wound healing

The signaling pathways that elicit monocyte migration are complex. Over the past few decades, extensive research has highlighted the critical role of neural-inflammatory interactions throughout all stages of wound healing, which has significant clinical implications (Wang et al. 2019).

The TM is a delicate tissue rich in sensory nerve fibers located near its outer surface. An acute perforation of the TM often results in an immediate and intense pain response in affected individuals. In this study, we hypothesize that the initial pain signal generated by tissue damage acts as a trigger for initiating an acute inflammatory response. This response includes vascular inflammation and the recruitment of monocytes. TRPV1 is recognized as a sensor for pain, cold, and itch (Ho et al. 2012; Marrone et al. 2017; Silverman et al. 2020). In other tissues, TRPV1-dependent pain sensation is known to rapidly provoke vascular inflammation, immune reactions, and the proliferation of precursor cells, accompanied by the swift release of neuropeptides such as Substance P (SP), neurokinin A, and calcitonin gene-related peptide (Du et al. 2019; Gazzieri et al. 2007; Tang et al. 2008; Oliveira et al. 2019; Corrigan et al. 2016). These neuropeptides significantly facilitate macrophage recruitment, vascular inflammation, and increased vascular permeability (Corrigan et al. 2016; Silverman et al. 2020; Foster et al. 2017; Breglio et al. 2020; Bujak et al. 2019; Li and Gupta 2019; Nidegawa-Saitoh et al. 2018). Additionally, a recent study demonstrated that cutaneous TRPV1+ neurons directly respond to noxious stimuli and are essential for innate immunity against various pathogens (Cohen et al. 2019).

In this study, TRPV1 was predominantly expressed in the trigeminal (TM) nerves and their associated trigeminal ganglia (Figure 6A and B). Further bulk RNA sequencing analysis of tissue from control and perforated wild-type (WT) animals at different time points indicated early signs of neuronal inflammation. Notably, we observed a significant upregulation of Tac1 expression in the perforated TMs. The Tac1 gene encodes four members of the tachykinin peptide family: Substance P, neurokinin A, neuropeptide K, and neuropeptide γ. This suggests that neural inflammation plays a role in the healing of TM wounds. By comparing monocyte recruitment and angiogenesis levels between WT and *Trpv1^-/-^* mice, we found that the *Trpv1^-/-^* mice exhibited significantly reduced monocyte recruitment and correspondingly lower angiogenic activity compared to the WT mice (Figure 7). Additionally, by day 7, the perforation in the Trpv1-/- mice had not fully sealed. Consistent with our findings, previous studies have shown that TRPV1 depletion impairs wound healing in both cutaneous and corneal tissues. (Ueno et al. 2023; Liu et al. 2022; Sumioka et al. 2014).

In conclusion, our data provide new insight into both the essential role of monocyte recruitment mediated by TRPV1 signaling and macrophage-associated angiogenesis in facilitating TM wound healing, as illustrated in Figure 8. These findings may open new translational opportunities for clinical intervention, as target of the TRPV1 signaling might enhance TM immunity or by promoting angiogenesis and boosting the blood circulation.

**Figure 8.**
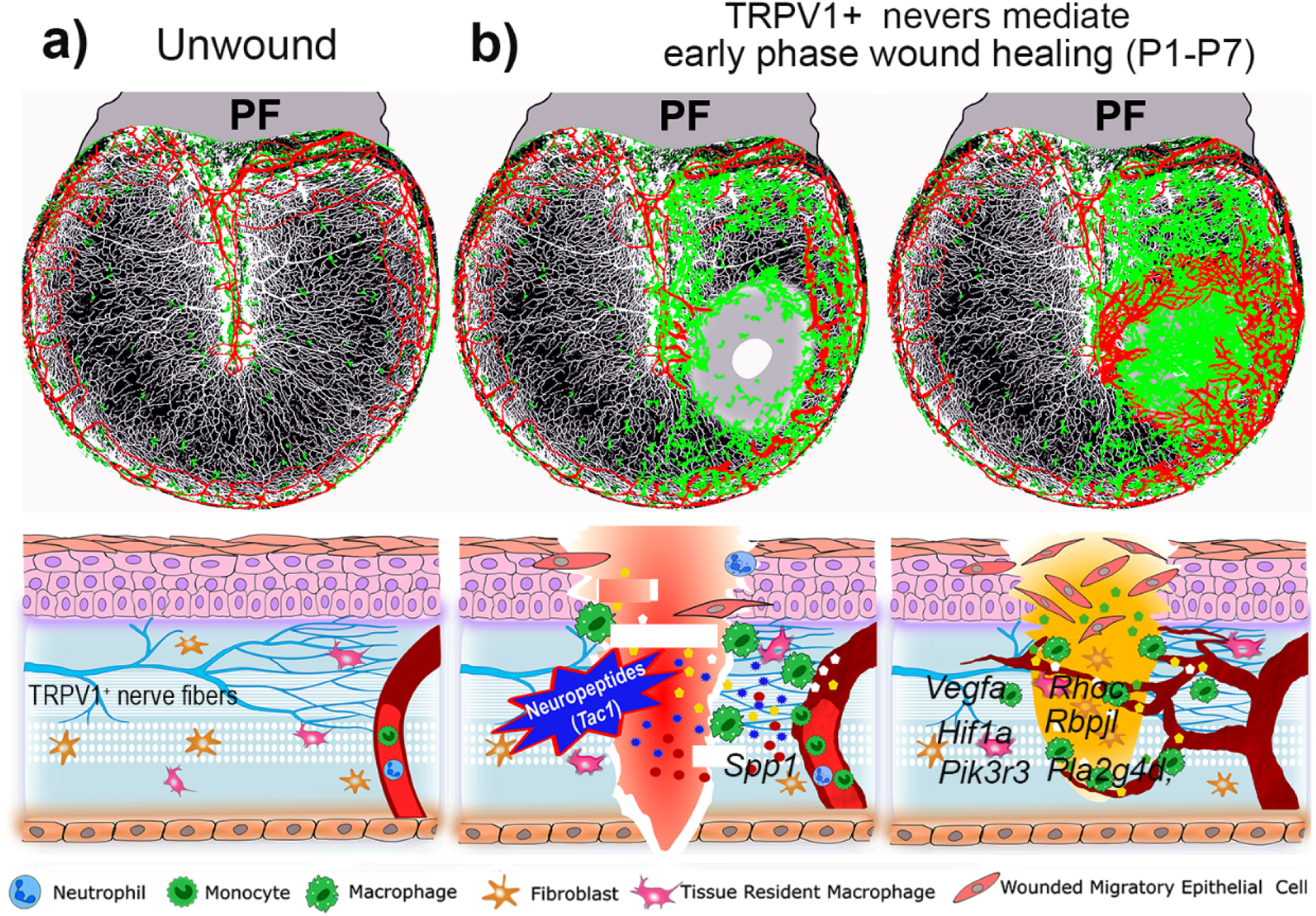
depicts the dynamics of monocyte migration and angiogenesis at various stages of TM wound healing. The upper section shows a whole mount view, while the lower section presents a cross-section. **a**) The figure illustrates the distribution patterns of TRPV1+ nerve fibers, blood vessels, and tissue-resident macrophages in normal TMs. **b**) On the left, it shows that damage to TRPV1+ nerve fibers results in the upregulation of the *Tac1* gene. This upregulation triggers the early release of neuropeptides, leading to vascular inflammation, monocyte recruitment, and communication between Spp1-mediated macrophages and endothelial cells following TM perforation. **b**) On the right, it details that increased monocyte recruitment and significant angiogenesis occur through various molecular signals, such as *Vegfa, Hif-1α,* and *Pla2g4d*, during the healing process of the TM wound.

## Acknowledgments

We would like to express our gratitude for the scRNA-seq dataset (GEO dataset: GSE196692) that was collected and shared with the scientific community by Dr. Aaron D. Tward’s research team.

## Author contributions

Yunpei, Zhang Ph.D. ── Formal analysis, Investigation, Methodology, Validation, Visualization, Writing – original draft and Editing.

Pingting Wang B.S. ── Formal analysis, Review and Editing

Lingling Neng M.D, Ph.D. ── Investigation, Visualization, Methodology

Kushal Sharma Ph.D. ── Methodology, Investigation, Validation, Visualization,

Allan Kachelmeier B.S. ── Writing –review and editing

Xiaorui Shi M.D., Ph.D ── Writing – Original draft, Review, Editing and Supervision

## Ethics

All animal experiments reported were approved by the Oregon Health & Science University Institutional Animal Care and Use Committee (IACUC IP00000968)

## Funding Information

This paper was supported by the following grants:

National Institute on Deafness and Other Communication Disorders R01 DC015781 to Xiaorui Shi.

National Institute on Deafness and Other Communication Disorders R01 DC010844 to Xiaorui Shi.

National Institute on Deafness and Other Communication Disorders R21 DC016157 to Xiaorui Shi.

National Institute on Deafness and Other Communication Disorders R01 DC022283 To Xiaorui Shi.

## Data availability

The datasets generated and/or analyzed during the current study are available from the corresponding author on reasonable request.

## Key resources table 1

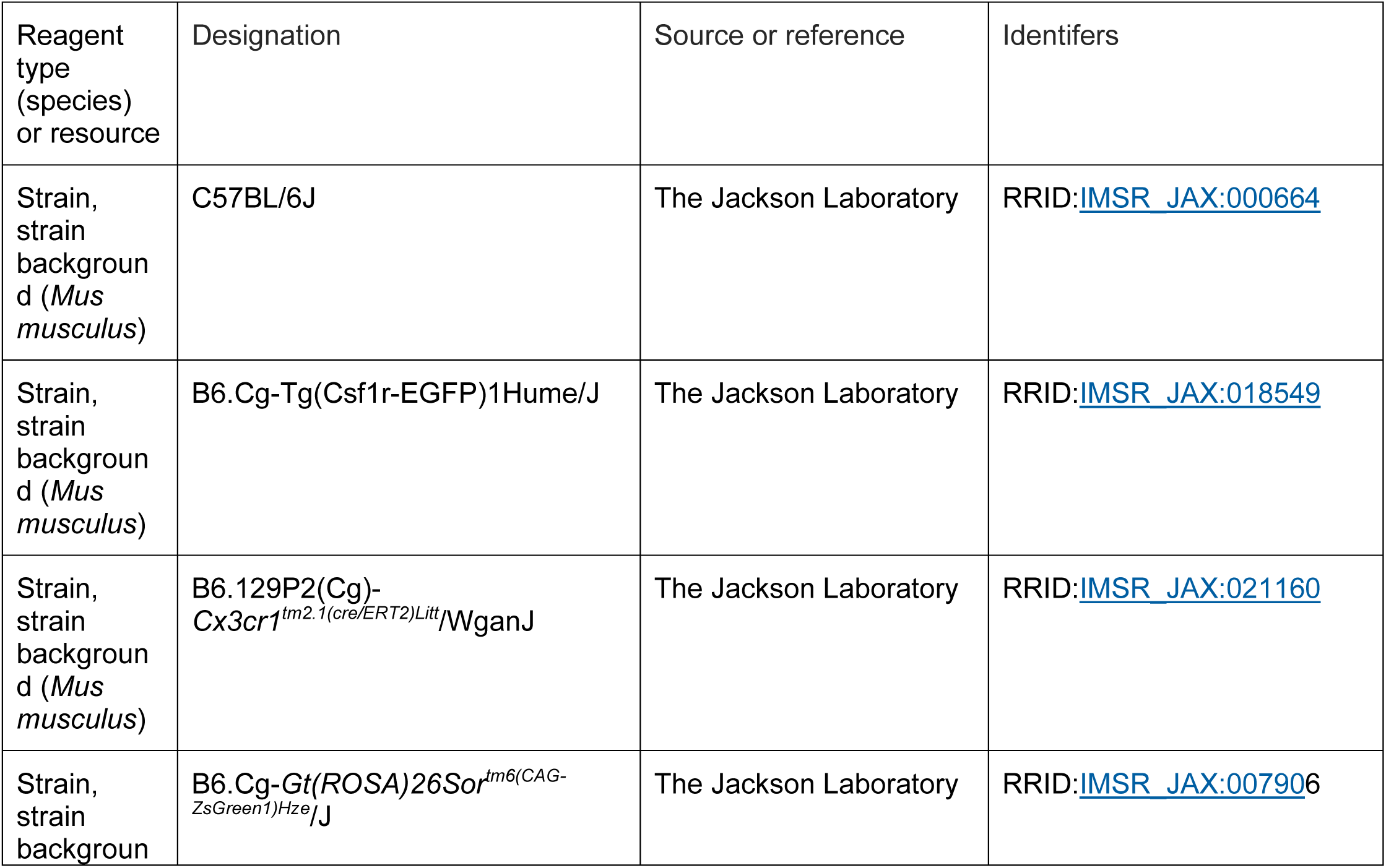
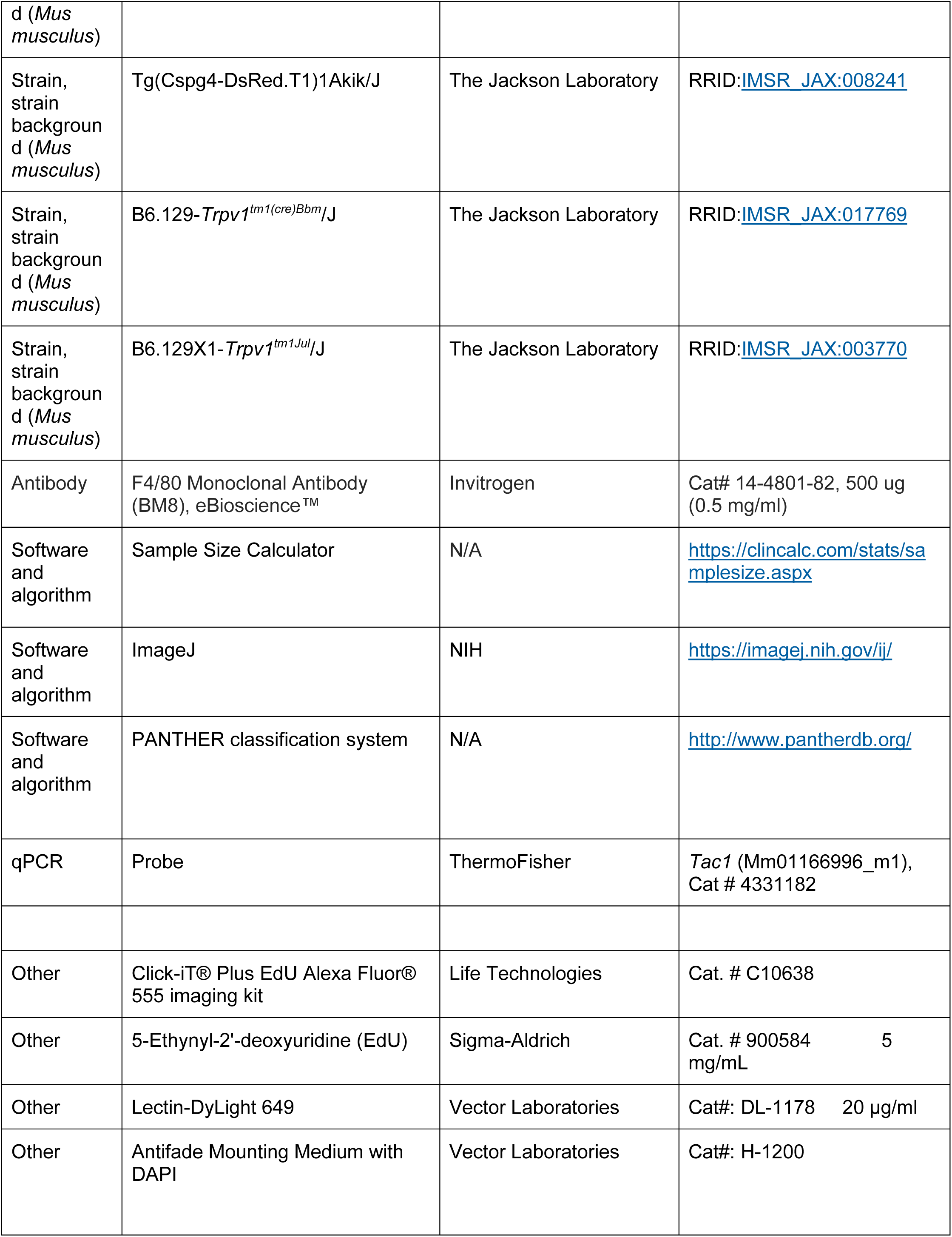

