## Supplemental Figure 1 for "Monocyte-derived macrophage recruitment mediated by TRPV1 is required for eardrum wound healing"

### Supplemental Materials

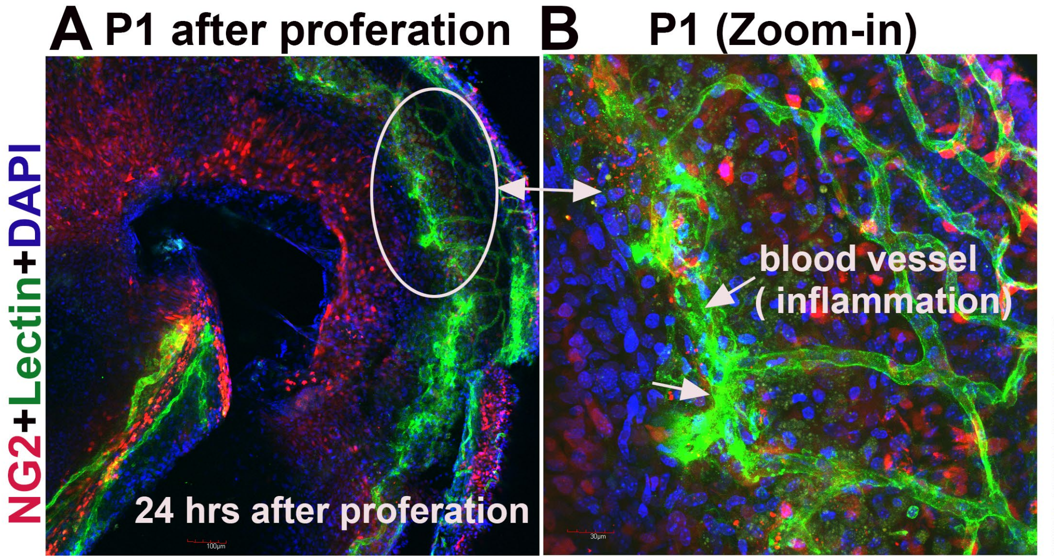


**Figure 1.** Inflamed vessels (arrows) in the vicinity of the wound region 24 hours after TM injury under low (**A**) and high (**B**) magnifications. The TM was isolated from a NG2+ fluorescence reporter mouse line 24 hours following TM perforation.
